# Occurrence of *Anoura geoffroyi* Gray, 1838, (Chiroptera: Phyllostomidae) in Quaternary deposits of Lagoa Santa, Brazil

**DOI:** 10.1101/2025.04.30.651513

**Authors:** Artur Chahud

## Abstract

Fossils of bats (Order Chiroptera) are considered rare due to the fragility of their bone structure, and are commonly represented only by fragments. In Brazil, most records correspond to extant species, with *Desmodus draculae* standing out as the only extinct species identified. The Lagoa Santa karst system is widely recognized for its paleontological and archaeological sites. Although many of its caves contain bat remains, detailed studies focusing on this group remain scarce. This study reports the first record of the species *Anoura geoffroyi* in Quaternary deposits. The skull was found in Cuvieri Cave, in a superficial layer associated with vertebrate remains currently present in the local fauna. Although the exact age of the specimen is uncertain, the presence of carbonate incrustations suggests that the skull remained exposed in the cave for an extended period before its final burial. Despite the impossibility of precisely defining the corresponding paleoenvironment, the current wide distribution of *Anoura geoffroyi* in the Cerrado Biome (Brazilian Savanna) suggests that the specimen may have lived in the region at a time when this biome was already predominant.

## INTRODUCTION

The Lagoa Santa region, located in eastern Brazil, stands out for its extensive and complex karst system, characterized by the presence of caves that house significant paleontological and archaeological remains. Scientific research in the area dates back to the 19th century, with the pioneering work of Peter Wilhelm Lund, who described both the extinct and contemporary fauna of the region (LUND, 1839; 1842). The fossil and subfossil assemblages of mammals from this region include a significant diversity of representatives of the extinct South American megafauna, in addition to species of the current fauna, including marsupials, xenarthrans, lagomorphs, rodents, ungulates and chiropterans. However, bats have historically been few studied, being mentioned sporadically in subsequent scientific literature.

The Order Chiroptera, which includes bats, is the second most diverse among current mammals and the only one with the ability to fly. The oldest fossil records of the group date back to the Eocene of Europe (TABUCE et al., 2019) and the United States of America (SIMMONS et al., 2008), when the species already presented anatomical characteristics similar to those of current taxa.

Despite a geological record dating back to the Paleogene, bat fossils are relatively rare due to the fragility of their anatomical structure, which makes their preservation in the paleontological record difficult (PAULA COUTO, 1979). They are often represented only by fragmented phalanges and metapods, and findings of jaws or skulls are uncommon.

In Brazilian caves, the remains of bats are relatively common and, in general, correspond to extant species (CZAPLEWSKI & CARTELLE, 1998; FRACASSO & SALLES, 2005). The only extinct species recorded to date in the country is *Desmodus draculae*, documented in deposits from the end of the Pleistocene.

In the Lagoa Santa region, fossil and subfossil remains of bats have been known since the 19th century, when WINGE (1893) identified bone elements attributed to current species of Chiroptera in local caves.

Among the most relevant paleontological sites in the region, Cuvieri Cave stands out, notable for the abundance of Quaternary osteological material and for the various taxonomic, taphonomic and sedimentological studies carried out at the site (HUBBE et al., 2011; HADDAD-MARTIM et al., 2017; CHAHUD, 2020, 2022; CHAHUD & OKUMURA, 2021b). However, no specific investigations have been conducted on the fossil remains of bats, which making this group the least explored among the mammals found in this cave.

Considering the scarcity of Chiroptera fossils and the lack of studies dedicated to the order in the region, the present work reports the discovery of a bat skull in deposits originally dated to the Pleistocene in Cuvieri Cave. The morphology of the specimen allows its attribution to the family Phyllostomidae, with affinities to the species Anoura geoffroyi (GRAY, 1838), representing the first record of this species in a Quaternary deposit in the Lagoa Santa region.

## MATERIAL AND METHODS

The Cuvieri Cave is located in the karst system of Lagoa Santa, Minas Gerais, and has two main passages: a larger one, currently obstructed, and a smaller one, approximately 1.5 m high by 1.0 m wide, which allows access to its interior. The cave contains three small vertical depressions, interpreted as natural traps, called *Locus* 1, 2 and 3 (Fig. 1), with respective depths of 16 meters, 4 meters and 8 meters (HUBBE et al., 2011; CHAHUD & OKUMURA, 2021a; 2023; CHAHUD et al., 2023).

**Figure 1.**
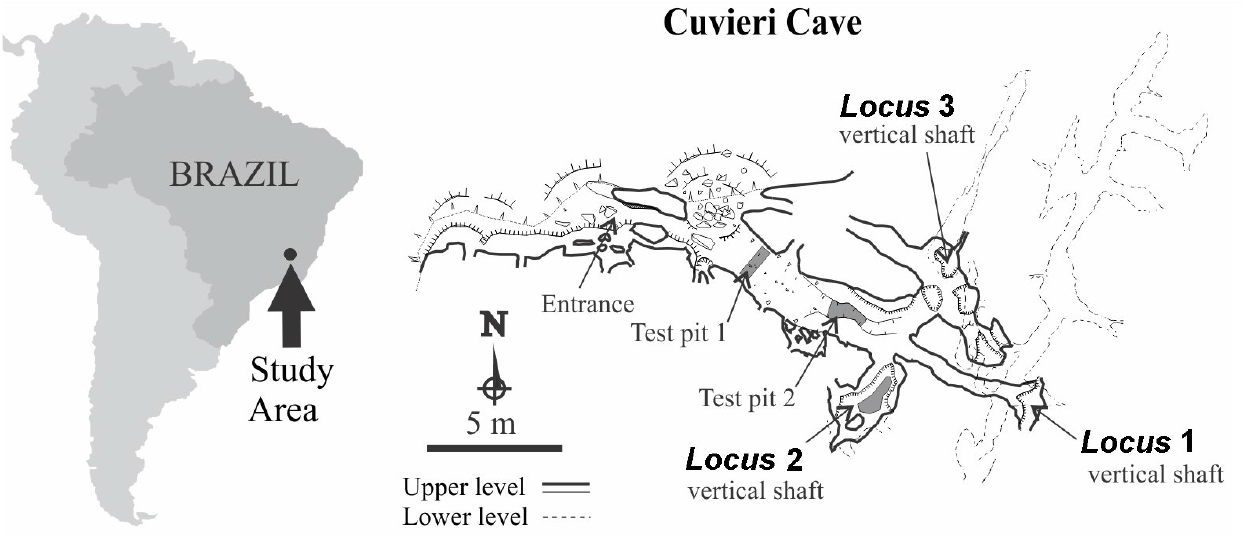
Geographic location of the study area and of Cuvieri Cave showing the position of *Loci* 1, 2 and 3 (map courtesy of Alex Hubbe and Grupo Bambuí for Speleological Research).

The specimen under study comes from *Locus* 3 and was identified in the most superficial layer of this cavity, placing it among the most recent materials recovered at the site. The dates available for *Locus* 3 indicate the presence of Pleistocene specimens even at superficial levels (MAYER et al., 2016; HADDAD-MARTIM et al., 2017). However, considering the organism’s flight capacity and its superficial stratigraphic position, it is not possible to state with certainty that the animal lived in the region during the Pleistocene. Therefore, a broader chronological assignment was chosen, located in the Quaternary.

The specimen was identified through direct observations and comparison with reference specimens, in addition to consulting specialized literature. The skull is duly catalogued and curated by the Laboratory of Human Evolutionary Studies at the Institute of Biosciences of the University of São Paulo (LEEH-IB-USP).

## SYSTEMATIC PALEONTOLOGY

Order Chiroptera Blumenbach, 1779

Suborder Microchiroptera Dobson, 1875

Family Phyllostomidae Gray, 1825

Subfamily Glossophaginae Bonaparte, 1845

Genus *Anoura* Gray, 1838

*Anoura geoffroyi* (Gray, 1838)

Figure 2

### Material

Almost complete skull, with few breaks and a small carbonate incrustation on the right side, belonging to an adult individual (CVL3-P1).

### Remarks

The skull (Figure 2) is small, with the left zygomatic arch partially preserved. The braincase tapers anteriorly, both on the anteroposterior axis and towards the base of the skull. There is no evidence of a sagittal crest, and the braincase is slightly rounded.

**Figure 2.**
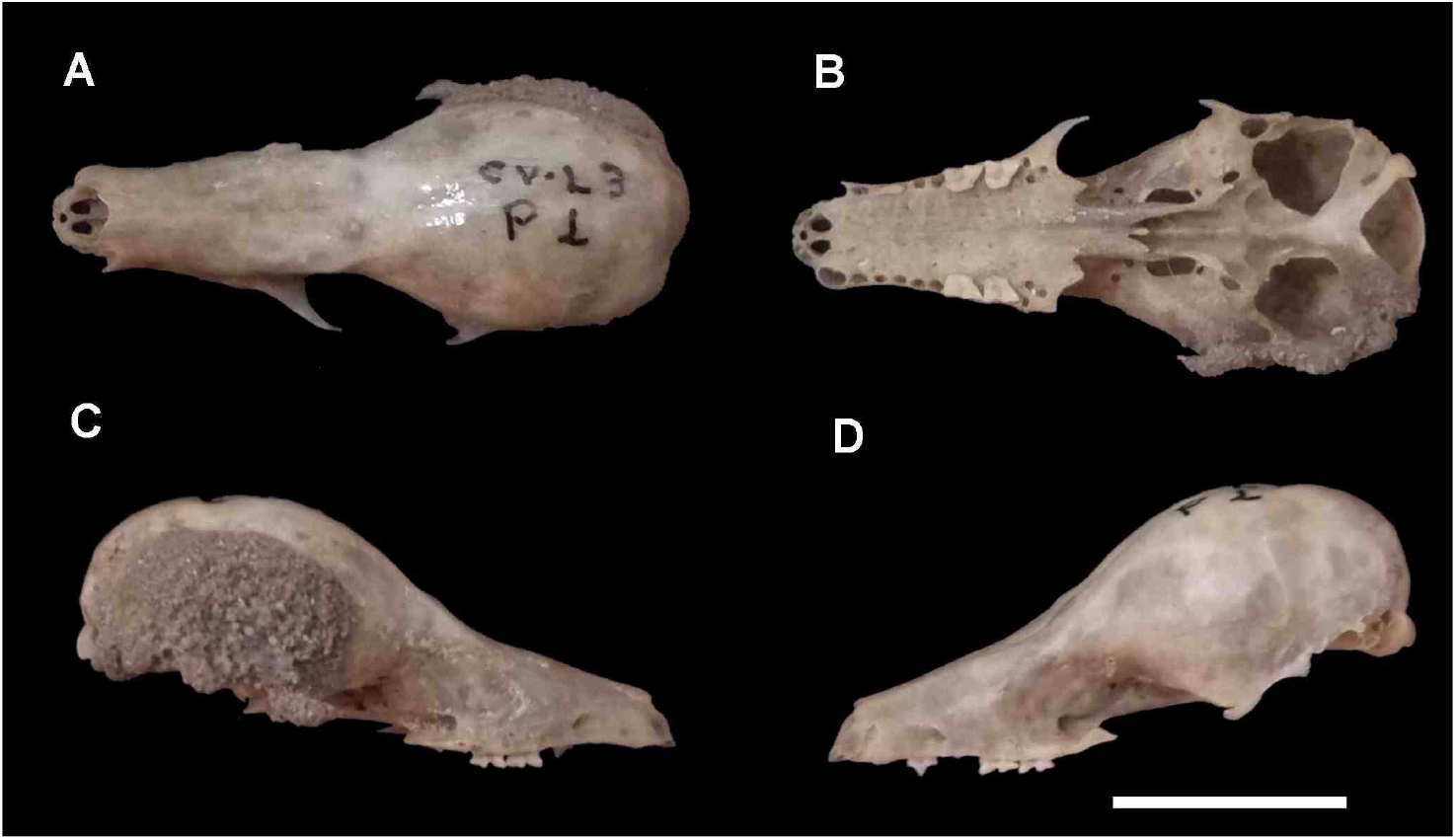
Skull of ***Anoura geoffroyi*** from Cuvieri Cave. A) Dorsal view; B) Ventral view; C) Right lateral view; D) Left lateral view. Scale: 10 mm.

The cranial morphology and the observed dental formula (I2/0, C1/1, P3/3, M3/3) are compatible with the characteristics of the genus Anoura. The specimen preserved a left premolar and the first and second molars on both sides of the maxilla. This dental condition distinguishes the specimen from other representatives of the Phyllostomidae family, which have more elongated skulls and may present variations in dental morphology or cranial conformation (FREEMAN, 1988; MORALES-MARTíNEZ & RAMíREZ-CHAVES, 2015; RAMíREZ-FRÁNCEL et al., 2020; BRANDÃO & HINGST-ZAHER, 2021).

The incisors are reduced and separated by a small median gap between the first incisors on each side. There is a slight spacing between the incisors and the canines, however, no diastemas are observed between the canines and the sequence of premolars and molars.

This individual is an adult, as evidenced by complete tooth eruption and development. The slight wear of the teeth suggests that it is a young adult.

### Discussion

The genus *Anoura* is represented in the Lagoa Santa region by two sympatric species: *Anoura geoffroyi* Gray, 1838, and *Anoura caudifer* Geoffroy, 1818. The skull of *A. geoffroyi* tends to be larger, more elongated and with a relatively thinner zygomatic arch compared to *A. caudifer* (ORTEGA & ALARCÓN-D, 2008; OPREA et al., 2009). Although the thickness of the zygomatic arch cannot be confirmed in the Cuvieri Cave specimen (Figure 2A and 2B), its general dimensions and the tapered morphology of the skull are compatible with those observed in specimens of *A. geoffroyi*.

The measurements presented in Table 1 are in accordance with the values recorded for *Anoura geoffroyi* by ORTEGA & ALARCÓN-D (2008) and are significantly higher than those observed for *A. caudifer* (OPREA et al., 2009), reinforcing the identification of the specimen as belonging to *A. geoffroyi*.

**Table 1.**
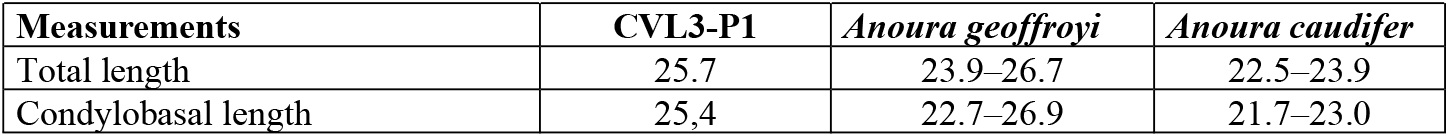
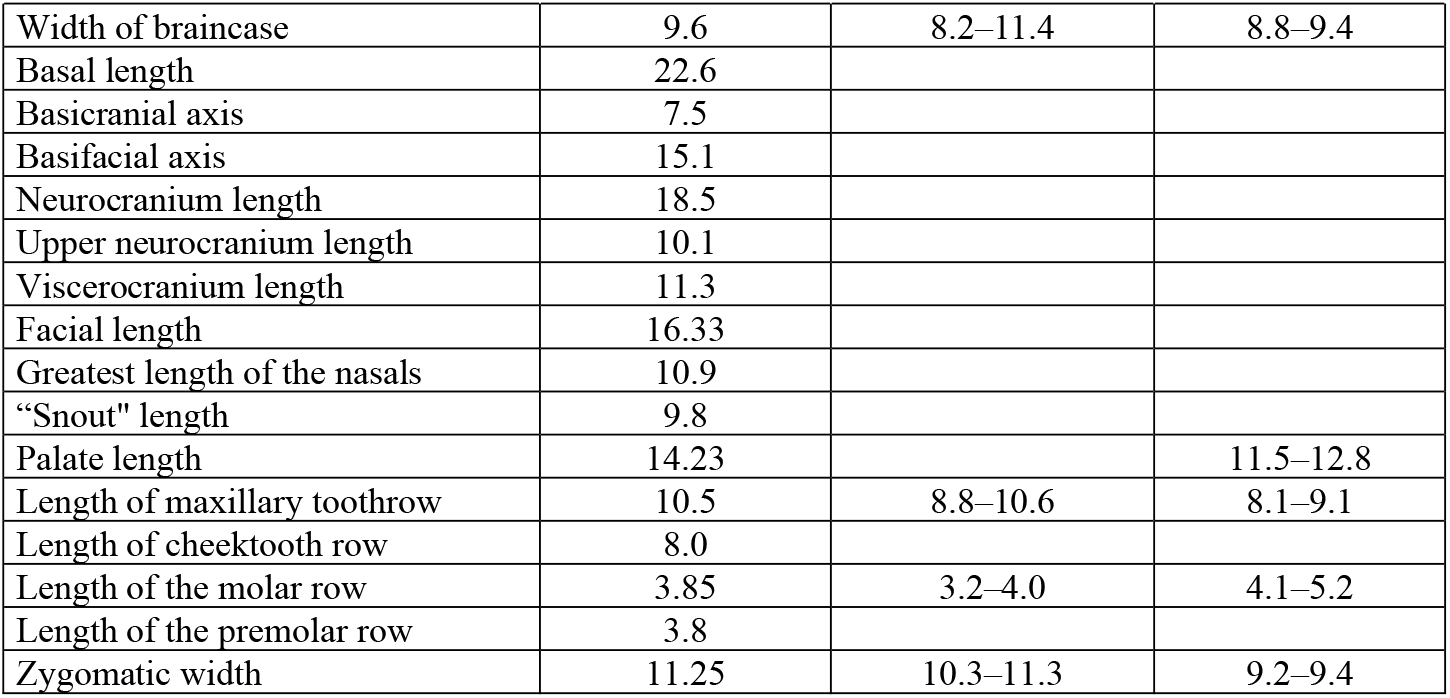
Measurements taken from the skull of *Anoura geoffroyi* from Cuvieri Cave. Measurements adapted from DRIESCH (1976). Ranges for *A. geoffroyi* adapted from ORTEGA & ALARCÓN-D. (2008), and for *A. caudifer* adapted from OPREA et al. (2009). All measurements are in millimeters (mm).

Comparison with specimens from the Atlantic Forest (BRANDÃO & HINGST-ZAHER, 2021) reveals no significant differences in dentition or cranial morphology compared to *A. geoffroyi*, supporting the taxonomic identification.

*Anoura geoffroyi* is widely distributed in the Neotropic, occurring from Mexico, northern and western South America and southeastern Brazil. This species has a generalist diet, consuming nectar, pollen, fruits and insects, with nectarivorous populations predominating in certain regions (McNAB, 1971; CABALLERO-MARTíNEZ et al., 2009).

In the Cerrado Biome, *Anoura geoffroyi* is one of the most frequent species among the Phyllostomidae, being recorded as the second most abundant in the region (ZORTÉA & ALHO, 2008). It is plausible that its occurrence was common in Lagoa Santa during the Quaternary. The specimen in question may have used the cave as shelter, a common behavior for individuals of the species.

## FINAL CONSIDERATIONS

The specimen was recovered from the surface layer of *Locus* 3 of Cuvieri Cave. Although these deposits are generally classified as Pleistocene based on dating of upper layers (MEYER et al., 2016; HADDAD-MARTIM et al., 2017; CHAHUD et al., 2023), the specimen in question comes from a layer with no evidence of extinct fauna or large vertebrates, presenting only remains of small vertebrates, such as rabbits and small rodents currently present in the Lagoa Santa region.

Considering that the specimen was found in the younger layers of the deposit and that it is a flying animal, there are no physical barriers restricting access to Locus 3 from the Pleistocene to the present. Therefore, it is not possible to assign a precise age. Although the exact chronology remains uncertain, the observed carbonate encrustations (Figure 2C) indicate that the specimen remained exposed in the cave for some time before its final burial. On the other hand, the fragility of the bones suggests that there was no significant remobilization.

Due to uncertainties regarding dating, it is not possible to determine a specific paleoenvironment for the time in which the specimen lived. However, Anoura geoffroyi is a species widely distributed in the Cerrado (Brazilian Savanna), making it plausible that the specimen inhabited the region during a period when this biome was already predominant, as it is today.

The fossil record of the genus *Anoura* dates back to a specimen of *A. caudifer*, identified by WINGE (1893), whose current classification was established by PAULA COUTO (1946). Similar to the specimen described here, the specimen of *A. caudifer* has also not been dated, and is widely considered to be from the Quaternary, but without ruling out the possibility that it is a recent specimen

## Acknowledgements

The author thanks the Dr. Maria Mercedes Martinez Okumura, responsible for LEEH (Laboratory for Human Evolutionary Studies), Department of Genetics and Evolutionary Biology, Institute of Biosciences of the University of São Paulo for permitting the preparation of the fossils in her laboratory.

## REFERENCES

Brandão M., Hingst-Zaher E. 2021. Atlas Craniano: mamíferos da mata atlântica e lista de espécies. São Paulo: Tijd Edições. 10.32673/9786588932018

Caballero-Martinez, L.A., Rivas-Apple, I.V. and Aguilera-Gomez, L.I. (2009). Feeding habits of Geoffroy’s tailless bat (Chiroptera: Phyllostomidae) in Ixtapan del Oro, Mexico State. Acta Zoologica Mexicana 25: 161–175. 10.21829/azm.2009.251609

Chahud, A., 2020. Occurrence of the sabretooth cat Smilodon (Felidae, Machairodontinae) in the Cuvieri cave, eastern Brazil. Palaeontologia Electronica. 23(2), a24. 10.26879/1056

Chahud A. 2022. Tafonomia de restos de Anura (Amphibia) do Holoceno da Gruta Cuvieri, estado de Minas Gerais, Brasil. Revista de Biologia Neotropical/Journal of Neotropical Biology, 19 (especial): 92–98. 10.5216/rbn.v19iesp.73433

Chahud, A. & Okumura, M. 2021a. The presence of Panthera onca Linnaeus 1758 (Felidae) in the Pleistocene of the region of Lagoa Santa, State of Minas Gerais, Brazil. Historical Biology, 33(10), 2496–2503. 10.1080/08912963.2020.1808975

Chahud, A. & Okumura, M. 2021b. The youngest Tapir of a Quaternary deposit of the Americas. Historical Biology, 33(10): 2400–2405. 10.1080/08912963.2020.1798420

Chahud, A.; Figueiredo, G. F.; Okumura, M. 2023. Cervidae and Tayassuidae of the Late Pleistocene from the Cuvieri Cave, eastern Brazil. Journal of South American Earth Sciences, 123: 104195. 10.1016/j.jsames.2023.104195

Chahud A., Okumura M. 2023. Cervidae and Tayassuidae from the Holocene deposits of the Cuvieri Cave, State of Minas Gerais, eastern Brazil; taxonomic and paleoenvironmental considerations. Historical Biology, 35: 74–83. 10.1080/08912963.2021.2022134

Czaplewski, N. J., & Cartelle, C. 1998. Pleistocene bats from cave deposits in Bahia, Brazil. Journal of Mammalogy, 79(3), 784–803. 10.2307/1383089

Driesch, V. D. A. 1976. A guide to the measurement of animal bones from archaeological sites: as developed by the Institut für Palaeoanatomie, Domestikationsforschung und Geschichte der Tiermedizin of the University of Munich (Vol. 1). Peabody Museum Press.

Fracasso, M. P. D. A., & Salles, L. D. O. 2005. Diversity of quaternary bats from Serra da Mesa (State of Goiás, Brazil). Zootaxa, 817(1), 1–19. 10.11646/zootaxa.817.1.1

Freeman, P. W. 1988. Frugivorous and animalivorous bats (Microchiroptera): dental and cranial adaptations. Biological Journal of the Linnean Society, 33(3), 249–272. 10.1111/j.1095-8312.1988.tb00811.x

Gray, J. E. 1838. A revision of the genera of bats (Vespertilionidae), and the description of some new genera and species. Magazine of Zoology and Botany 2:483–505.

Haddad-Martim, P. M., A. Hubbe, P. C. F. Giannini, A. S. Auler, L. B. Piló, M. Hubbe, E. Mayer, X. Wang, H. Cheng, R. L. Edwards & W. A. Neves, 2017. Quaternary depositional facies in cave entrances and their relation to landscape evolution: The example of Cuvieri Cave, eastern Brazil. Catena 157: 372–387. 10.1016/j.catena.2017.05.029

Hubbe, A.; Haddad-Martim, P.M.; Hubbe, M.; Mayer, E.L.; Strauss, A.; Auler, A.S.; Pilo, L.B.; Neves, W.A. 2011. Identification and importance of critical depositional gaps in pitfall cave environments: the fossiliferous deposit of Cuvieri Cave, eastern Brazil. Palaeogeography Palaeoclimatology Palaeoecology, v.312, p.66–78. 10.1016/j.palaeo.2011.09.010

Lund, P.W. 1839. Coup d’oeil sur les espèces éteintes de Mammifères du Brésil, extrait de quelques mémoires présentés à l’Académie Royale des Sciences de Copenhague. Annales des Sciences Naturelles, 11:214–234.

Lund, P.W. 1842. Blik paa Brasiliens Dyreverden för Sidste Jordomvaeltning. Fjerde Afhandling: Fortsaettelse af Pattedyrene. Lagoa Santa, Det Kongelige Danske VidenskabernesSelskabs Naturvidenskabelige og Mathematike Afhandlinger, 9:137–208.

Mayer E.L.; Hubbe A.; Kerber L.; Haddad-Martim P.; Neves W. 2016. Taxonomic, biogeographic, and taphonomic reassessment of a large extinct species of paca from the Pleistocene of Brazil. Acta Palaeontologica Polonica, 61(4):743–758. 10.4202/app.00236.2015

Mcnab, B. K. 1971. The structure of tropical bat faunas. Ecology 52: 352–358. 10.2307/1934596

Morales-Martínez, D. M., & Ramírez-Chaves, H. E. 2015. The distribution of bats of genus Lasiurus (Vespertilionidae) in Colombia, with notes on taxonomy, morphology and ecology. Caldasia, 37(2), 397–408. 10.15446/caldasia.v37n2.54392

Oprea, M., Aguliar, L., & Wilson, D. E. (2009). Anoura caudifer (Chiroptera: Phyllostomidae). Mammalian Species, (844), 1–8. 10.1644/844.1

Ortega, J., & Alarcón-D, I. (2008). Anoura geoffroyi (Chiroptera: Phyllostomidae). Mammalian Species, (818), 1–7. 10.1644/818.1

Paula Couto, C. 1946. Atualizando da nomenclatura generica e especifica usada por Herluf Winge, em “E Museo Lundii.” Estudos Brasileiros de Geologia, 1:59–80.

Paula Couto, C. 1979. Tratado de Paleomastozoologia. Academia Brasileira de Ciências, Rio de Janeiro.

Ramirez-Fráncel, L. A., García-Herrera, L. V., & Reinoso-Florez, G. 2020. Using MaxEnt modeling to predict the potential distribution of Platyrrhinus ismaeli (Phyllostomidae). Therya, 11(2), 203–212. 10.12933/therya-20-843

Simmons, N. B., Seymour, K. L., Habersetzer, J., & Gunnell, G. F. 2008. Primitive Early Eocene bat from Wyoming and the evolution of flight and echolocation. Nature, 451(7180), 818–821. 10.1038/nature06549

Tabuce, R.; Antunes, M. T.; Sigé, B. 2009. A new primitive bat from the earliest Eocene of Europe. Journal of Vertebrate Paleontology. 29 (2): 627–630. 10.1671/039.029.0204

Winge, H. 1893. Jordfundne og nulevende Flagermus (Chiroptera) fra Lagoa Santa, Minas Geraes, Brasilien: med udsigt over Flagermusenes indbyrdes Slaegstkab. E Museo Lundii, 2:1–92. 10.5962/bhl.title.14824

Zortéa M., & Alho. C. J. R. 2008. Bat diversity of Cerrado habitat in central Brazil Biodiversity and Conservation. 10.1007/s10531-008-9318-3

